# Temporal microstructure of dyadic social behavior during relationship formation in mice

**DOI:** 10.1101/711036

**Authors:** Won Lee, Jiayi Fu, Neal Bouwman, Pam Farago, James P. Curley

## Abstract

Understanding the temporal dynamics of how unfamiliar animals establish dominant-subordinate relationships and learn to modify their behavior in response to their social partner in context-appropriate manners is critical in biomedical research concerning social competence. Here we observe and analyze the microstructure of social and non-social behaviors as 21 pairs of outbred CD-1 male mice (*Mus Musculus*) establish dominant-subordinate relationships during daily 20-minute interaction for five consecutive days. Using Kleinberg burst detection algorithm, we demonstrate aggressive and subordinate interactions occur in bursting patterns followed by quiescence period rather than in uniformly distributed across social interactions. Further, we identify three phases of dominant-subordinate relationship development (pre-, middle-, and post-resolution) by combining phi-coefficient and difference methods used to determine at which bursting event mice resolve dominant-subordinate relationships. Using First Order Markov Chains within individuals we show dominant and subordinate animals establish significantly different behavioral repertoire once they resolve the relationships. In both dominant and subordinate mice, the transitions between investigative and agonistic behavior states are not common. Lastly, we introduce Forward Spike Time Tiling Coefficient, the strength of association between the given behavior of one individual with the target behavior of the other individual within a specified time window. With this method, we describe the likelihood of a mouse responding to a behavior with another behavior differ in pre- and post-resolution phases. The data suggest that subordinate mice learn to exhibit subordinate behavior in response to dominant partner’s behaviors while dominant mice become less likely to show subordinate behaviors in response to their partners’ action. Overall, with the tool we present in this study, the data suggest CD-1 male mice are able to establish dominance relationships and modify their behaviors even to the same social cues under different social contexts competently.

## Introduction

A significant challenge in understanding how the brain regulates dynamic changes in social interactions is the analysis of the microstructure of behavior. In studies of dyadic social interaction in laboratory animals such as mice, most research has focused on analyzing the duration and frequency of, or latency to perform, social behaviors in single short duration tests. These outcome measures are commonly assessed independently in each individual without necessarily determining how each animal’s behavior is influencing, or is influenced by, the social partner. While these endpoints do provide useful information as to the overall behavior of individual mice, they do not give insight into the temporal changes that occur as animals engage in social interactions. Across taxa, one particularly significant social relationship is the establishment of stable dominant-subordinate social roles over time by engaging in repeated agonistic contests. Once these relationships are formed, dominant and subordinate animals differ in their physiology, neurobiology, and behavior [1,2]. The ability of animals to learn how to dynamically change their social behavior with respect to another individual is termed social competence [3]. The behavioral dynamics through which dyads go from having no previous interactions to showing stable individual differences in agonistic and defensive behavior appear to vary substantially across species and contexts [4–6]. Understanding precisely how animals change their response to other individuals’ behaviors over time during the establishment of such relationships will help identify individuals who exhibit elevated or compromised social competence, and will ultimately facilitate understanding the neural mechanisms of social interaction.

A number of methods have been proposed for studying the temporal dynamics of social interaction in laboratory animals but these are still underutilized. One approach has been to analyze the First Order Markov transitions between successive behaviors produced by one individual to identify those behaviors that transition between each other more frequently than expected by chance. This method has been used to analyze non-social rodent behaviors exhibited in a solitary environment such as the forced swim test, elevated plus maze, open field, and exploration box [7–11] as well as the sequential pattern of social behaviors made by one individual in the resident-intruder paradigm and during social play [12–19]. An advantage of this method is that it can reveal contingencies occurring between two behaviors made by each individual, but it is also limited in that it can underestimate the association of two behaviors if they are interrupted by other behaviors. Furthermore, this type of analysis does not address interactions between behaviors made separately by each individual. To address this issue, it is possible to synchronize sequences of each subject to create one overall sequence for event-lag based sequential analyses [20–22], or utilize sequential timed-window approaches that facilitate the identification of behaviors made by one individual that occur within a window of the onset of another behavior made by their partner [23–25].

In the present study, we paired outbred CD-1 male mice with each other for twenty minutes per day for five days. We first sought to characterize precisely how and when each dyad successfully resolved their relative social status with one male becoming dominant and the other subordinate. Prior studies have defined dominant males as those that continue to aggress while subordinate males are those that consistently yield to the dominant [26]. However, this relatively simple definition lacks clarity as how to computationally determine the exact time-point at which this consistent pattern of social interaction becomes entrenched. Here we introduce the Kleinberg burst detection algorithm to characterize the bursting patterning of aggressive and subordinate interactions in pairs and present two methods to identify at which burst the social status of each animal is fully resolved. Further, using First Order Markov Chains within individuals we identify changes in the microstructure of social behavior from before to after this relationship resolution. We also introduce a time-based pairwise correlation method (based on a method previously used to calculate the correlation of neuronal spikes in electrophysiological time-series data [27]) for identifying significant associations between the given behavior of one individual with the target behavior of another individual within a specific time window. Understanding how the microstructure of these dyadic interactions changes over time is critical to identifying how behaviors may be initiated or inhibited in response to identical behavioral cues under different social contexts and how animals function in a socially competent manner.

## Materials and Methods

### Subject animals, husbandry and behavioral testing

A total of forty-two male outbred CD-1 males (*Mus Musculus*) aged 8 weeks were used in the study. We chose to use male subjects as males reliably form dominant-subordinate roles in dyads whereas females do not. All animals were bred in the animal facility in the Department of Psychology at Columbia University, housed with constant temperature (21–24°C), humidity (30-50%) and a 12/12 light/dark cycle with white light (light cycle) on at 2400 hours and red lights (dark cycle) on at 1200 hours. All animals were housed post-weaning with littermates in groups of 3 in standard sized cages (27 × 17 × 12 cm). At the beginning of the study male mice were singly housed for 7 days. On Day 8 of isolation, animals were habituated to a social interaction testing chamber (31.75 cm × 27.3 cm) by placing them alone for 20 minutes in the chamber. The following day, males were placed in pairs into the social interaction testing chamber for 20 minutes. Following 20 minutes of behavioral testing, animals were separated and placed back into their home cages. For four more successive days animals were paired with the same individual on each day, resulting in a total of five days of behavioral testing. All animals were tested during the first five hours of the dark phase of the light cycle. Each animal was identified by a black ink mark on its tail. Each pair of males was from a separate litter and had had no prior social experience with each other. Throughout all housing and testing periods, animals were provided with pine shaving bedding. Animals were also housed with nestlets to provide cage enrichment. All procedures were approved and conducted in accordance with the Columbia University Institutional Animal Care and Use Committee and National Health Institute (NIH; Bethesda, MD, USA) animal care guidelines. Behaviors were coded from videos using Observer XT (Noldus V11.5) by trained observers. We recorded the onset, termination and duration of every behavior exhibited by each mouse during each social interaction test. Our behavioral ethogram and categorization of behaviors into different states is shown in **Table 1** and was based on those previously used for mouse social interactions [12]. There are four different behavior states of social behaviors: Aggressive, subordinate, investigative, and other social behaviors states. Throughout this manuscript, we refer to the other social behavior state as social behavior state. In total across the 21 pairs tested on five separate days we recorded a total of 105,325 behavioral events.

**Table 1.** Behavioral ethogram and categorization of behaviors.

| State | Behavior | Description |
| --- | --- | --- |
| **Aggressive Behaviors** | Lunge | Mouse lunges an attack at partner with or without making contact to partner |
|  | Bite | Mouse bites partner on any part of body |
|  | tail rattle | Mouse rattles their rigid tail rapidly |
| **Subordinate Behaviors** | Flee | Mouse moves rapidly to create maximal distance between themselves and partner |
|  | Freeze | Mouse stays completely immobile in presence of partner typically also showing piloerection |
|  | subordinate posture | Mouse rears head and exposes nape and flank to partner while submitting |
| **Investigative Behaviors** | sniff-head | Mouse olfactory investigates the head of partner with or without direct contact |
|  | sniff-body | Mouse olfactory investigates the body of partner with or without direct contact |
|  | sniff-anogenital | Mouse olfactory investigates the anogenital region of partner with or without direct contact |
|  | sniff-follow | Mouse olfactory investigates the partner whilst partner is moving away from them |
| **(Other) Social Behaviors** | Pursue | Mouse follows the partners whilst there are moving without sniffing them |
|  | Allogroom | Mouse grooms partner on their fur on any part of their body with mouth or forepaws |
|  | side-by-side contact | Mouse sits in contact with the partner without freezing or engaging in other social behaviors |
| **Repetitive Behaviors** | digging | Mouse digs and moves bedding around the cage with their mouth or forepaws |
|  | self-grooming | Mouse grooms their own fur with mouth and forepaws |
| **Activity** | active moving | Mouse is engaged in locomotor activity around the cage without exhibiting any other behaviors |
|  | rearing | Mouse rises up on its hind legs and stretches the body |
|  | jumping | Mouse leaps into the air |
| **Inactivity** | idle | Mouse does not engage in any behaviors but is immobile |

### Statistical analysis

#### Differences in behaviors of dominant and subordinate individuals over days

All Bayesian generalized linear mixed effect models in the study were analyzed using the brms R package [28] with default priors in the package. We visually inspected Markov Chain Monte Carlo (MCMC) chains for proper mixing and used standard MCMC convergence diagnostics [29] for each model. To test for differences in the duration of each behavior between dominant and subordinate animals across days we used Bayesian fitted in Stan [30] via with the duration of behavior as the outcome variable, individual social status (dominant vs subordinate) and day as fixed factors and pair-id and individual mouse-id as random factors. We fitted data with either Gaussian distribution (sniff-head, sniff-body, side-by-side contact) or lognormal distribution (aggressive behavior state, subordinate behavior state, anogenital sniffing, sniff-follow, allogroom).

#### Burst detection

We identified bursts in aggressive and subordinate behavior states using Kleinberg’s burst detection method [31]. Briefly, for each dyad on each day we generated a sequence of the times of all aggressive and subordinate behaviors exhibited by either individual. Kleinberg’s algorithm then identifies bursts of activity using an infinite hidden Markov model to detect periods of increased activity in the time-series of discrete events (see Kleinberg [31] for more details). We used the R package ‘bursts’ for analysis [32]. In the model we set gamma as 0.3 and the second order bursts were determined as the bursts. Change in burst frequency over 5 days was assessed with zero inflated Poisson family in the brms R package. Change in burst durations over days was fitted with a lognormal distribution. The effect of days on the proportion of aggressive and subordinate behaviors transpired within bursts was tested using logistic regression. Pair-ids were added to each model as a random factor.

#### Identifying resolved dominant-subordinate relationships across bursts

For every burst of each dyad, a 2×2 matrix was constructed with the four cells being a,b,c,d, where *a* = frequency of aggressive behavior by the dominant male, *b* = frequency of subordinate behavior by the dominant male, *c* = frequency of aggressive behavior by the subordinate male, and *d* = frequency of subordinate behavior by the subordinate male. Animals that showed a higher proportion of aggressive behavior and lower proportion of subordinate behavior in the last two bursts were considered as the eventual dominant animals. Two methods were used to identify the burst at which each dyad resolved their dominant-subordinate relationship.

##### i) Phi-Coefficient Method

The phi-coefficient is calculated by 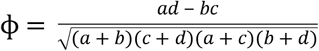 [33]. ϕ ranges between −1 and +1. +1 indicates that the dominant male is exclusively aggressive and shows no subordinate behavior (b=0) and that the subordinate male shows only subordinate behavior and no aggressive behavior (c=0). A value of ϕ=0 indicates that both animals show equivalent amounts of both behaviors (a/c=b/d). The significance of each ϕ coefficient was determined by calculating the chi-square value using the formula *X*^2^ = *N*ϕ^2^ where *N* = *a* + *b* + *c* + *d*. For every burst the phi-coefficient and its associated p-value was calculated. This is a very stringent method as it requires a sufficient number of behaviors to occur within each burst and for them to occur primarily on the off-diagonal of the matrix (i.e. one animal almost exclusively is aggressive, and the other animal is almost exclusively subordinate) to show a significant effect. This method also requires both animals to exhibit behaviors as it is not possible to calculate the phi-coefficient for bursts where one partner show neither aggressive nor subordinate behavior. Therefore, we excluded any bursts with less than 6 behavior instances. Relationships were determined to be resolved at the first burst following which all remaining bursts had ϕ > 0 and p<0.1. If there were fewer than 3 bursts left, relationships were adjudged to be resolved if the final or final two bursts were also ϕ > 0 and p<0.1. If no bursts satisfied these criteria, then the relationship was determined to be unresolved. The Wilcoxon Signed Rank Test was used to test if the phi-coefficient of the very first burst of each dyad was different from 0. The exact binomial test was used to test the significance of the success rate in dominant-subordinate resolution.

##### ii) Difference Method

Dyads were adjudged to be resolved when the difference of aggressive behaviors between the dominant and the subordinate was larger than that of the difference in subordinate behaviors between both males: (*a* – *c*) ≥ (*b* – *d*). Relationships were determined to be resolved at the first burst following which all remaining bursts satisfied this criteria.

#### Behavioral changes across pre-resolution, middle, post-resolution phases

For each dyad, a relationship was considered ‘pre-resolution’ for the time prior to the burst when it was first considered resolved according to the difference method; a relationship was considered ‘fully resolved’ for the time following the burst when it was considered to be resolved according to the phi-coefficient method; a relationship was considered to be in the ‘middle’ for the time in between these two time points. These criteria ensured that no relationship was considered ‘pre-resolution’ when we had at least some evidence to the contrary using the less-stringent difference methods, and no relationship was considered ‘post-resolution’ until there was strong evidence from the stringent phi-coefficient method. For each dyad the duration of behavior exhibited by the eventual dominant and subordinate male during each phase (pre-resolution, middle, post-resolution) for every behavior was calculated. From this, the proportion of time (ranging between 0 and 1) that each animal engaged in each behavior was calculated. Significant differences in duration of each behaviors between dominant and subordinate males at each phase were tested using paired Wilcoxon-Signed Rank Sum Tests.

#### First Order Markov Chains analysis within individuals

Unique non-repeating sequences were generated for each subject’s behavior during the pre- and post-resolution phases. Simply, for each subject, each sequence contained the order of behavioral transitions with no repeats allowed. For each sequence a frequency matrix of observed first-order transitions was generated. Only one dyad (L) for the pre-resolution phase was excluded from the analysis as the sequence of behaviors prior to resolution was too short. To test for non-uniformity in transitions in each matrix, we applied likelihood ratio tests [34]. For each matrix, we calculated the likelihood ratio statistic (G^2^) by

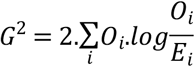

where *O* = the observed count of transitions, *E* = the expected count of transitions, *i* = each cell of the matrix. As no behavior can repeat itself each matrix has structural zeros along its diagonal, therefore we used iterative proportional fitting using the ‘mipfp’ R package to generate the expected counts of transitions and adjusted degrees of freedom accordingly [34,35]. To identify behavioral transitions that occurred at a frequency significantly higher than expected by chance, we used permutation tests. Each behavioral sequence was randomly permuted 1000 times and we ensured that no behavior could repeat itself to maintain the key structural property of original sequences [36]. P-values for each behavioral transition in each sequence were calculated by dividing the number of times each first-order transition occurred more frequently in the permuted sequences than in the observed sequence by the number of permutations. Transitions were conservatively considered to occur significantly more frequently than expected by chance within groups (dominant pre-resolution, dominant post-resolution, subordinate pre-resolution, subordinate post-resolution) if the median p-value over all individuals within that group across dyads was p<=0.01. For all transitions a transition probability was calculated. This ranges from 0-1 and represents how likely a behavioral transition selected at random would be that particular transition. For each group we also calculated the median transition probability of each behavioral transition by taking the median value across all individuals in each group. As many behavioral transitions occur at very low frequencies, we considered behavioral transitions that occur with a median transition probability of at least 0.075 to be meaningful.

#### Timed-window cross-correlation analysis

For each individual in each pair a time-series of the onset times of each behavior was generated. To calculate the cross-correlations between a given behavior of individual A and a target behavior of individual B, we adapted the Spike Time Tiling Coefficient (STTC) method developed by Cutts & Eglen [27]. The STTC method has numerous advantages over other cross-correlation methods in that it does not require assumptions about the frequency or distribution of events being correlated in each time-series but accounts for the overall frequency of each behavior in the calculation of the coefficient. In brief, we adapted the STTC formula to consider how likely a target behavior of an individual was to occur following a given behavior of the other individual within a 2 second window. For instance, suppose there are two times series *A* and *B. A* represents the onset times of the given behavior of individual A and *B* represents the onset times of the target behavior of individual B. We calculate the Forward STTC (FSTTC) of *A*→*B* as follows:

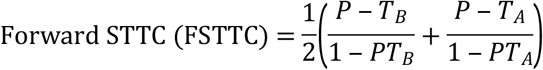

Where *P* is the proportion of behavioral onsets from *B* that lie within time Δ*t* after any behavioral onset from *A. T_A_* is the proportion of total observation time which lies within time Δ*t* after any spike from *A. T_B_* is the proportion of total observation time which lies within time Δ*t* after any spike from *B*. We used Δ*t* = 2*s* throughout. The algorithm was written in C++ with a ready application in R. We calculated FSTTC values among behavior states (see **Table 1**) and behaviors of each dyad in both directions (DOM➔SUB, SUB➔DOM) across each relationship phase (pre- and post-resolution). Median differences between the directions in both phases were tested using paired Wilcoxon Signed Rank Tests.

## Results

### Changes in duration of behaviors of dominants and subordinates over days

We evaluated how each individual changed their behavior across the 5 days of social interaction. We identified individuals as dominant or subordinate based on which animal exhibited clear aggressive behavior without yielding by the end of Day 5 (see **Methods**). **Fig 1** shows the total duration of all aggressive and subordinate behavior states across 5 days for the dominant and subordinate animal from each dyad. As clearly shown in **Fig 1**, dominant individuals engage in significantly more aggressive behavior than subordinate males (β_dom-sub_= 0.39, 95% credibility interval = [0.16, 0.62]). Both dominants and subordinates decreased the time they engaged in aggressive behaviors over days (β_days_ = −0.32 [−0.40, −0.24]). Likewise, dominant individuals exhibited significantly less subordinate behavior compared to subordinates (β_dom-sub_= −1.72 [−2.57, −1.12]) and overall subordinate behavior exhibited by both dominants and subordinates also significantly declined over time (β_days_= −0.08 [−0.15, −0.01]). **Supplementary Fig S1** shows the plots for the daily durations of investigative and other social behaviors exhibited by each mouse in each dyad. Generally, dominant mice engaged in grooming subordinate mice significantly more frequently than vice versa (allogroom: β_dom-sub_= 23.97 [10.18, 37.45]). Conversely, subordinates engaged significantly more frequently in anogenital sniffing (β_dom-sub_= −0.17 [−0.36, −0.00]) and side-by-side contact (β_dom-sub_= −48.35 [−66.41, −30.64]). There were no significant differences in sniff-follow, sniff-body and sniff-head between dominants and subordinates (sniff-follow: β_dom-sub_= −0.19 [−0.82, 0.39]; sniff-body: β_dom-sub_= −7.33 [−22.03, 7.69]; sniff-head: β_dom-sub_= 18.02 [−4.29, 40.90]). Across the five days, both individuals engaged in anogenital sniffing (β_days_= −0.14 [−0.19, −0.09]), sniff-follow (β_days_= −0.83 [−1.04, −0.66]), and allogrooming (β_days_= −6.01 [−8.88, −3.18]) less frequently over time. Conversely, mice increased time engaging in sniff-head (β_days_= 24.42 [18.91, 29.94]) and side-by-side contact (β_days_= 8.87 [4.22, 13.51]). The time mice engaged in sniff-body did not change over 5 days (β_days_= 0.13 [−3.37, 3.47]).

**Fig 1.**
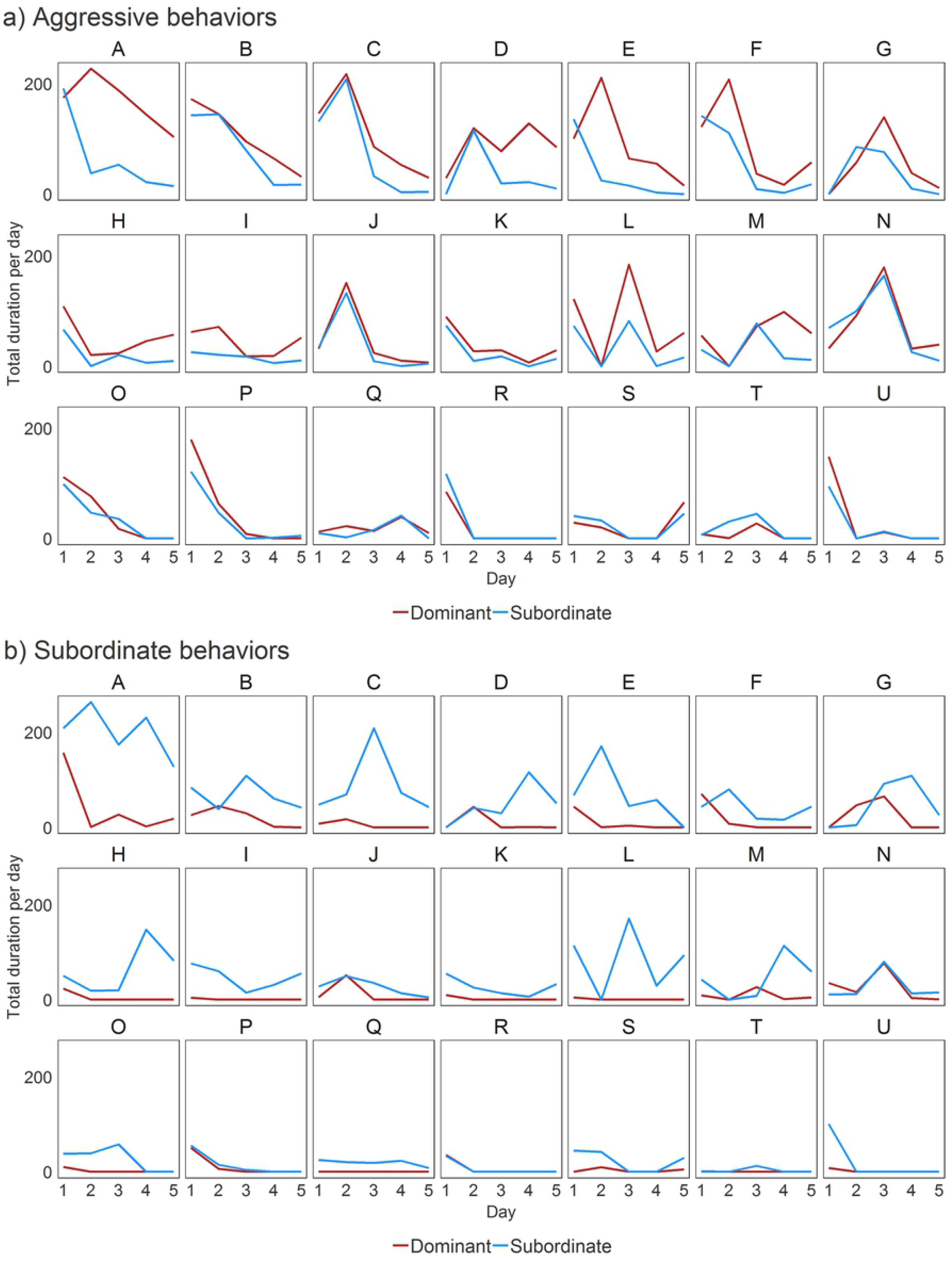
Individual differences in duration of aggressive and subordinate behaviors across days. Dyads (A-U) are listed in rank order of total aggression. Red and blue lines represent animals determined to be dominant and subordinate in each dyad by the end of Day 5.

### Temporal patterning of behavior over time

In this section, we describe the variation across dyads in the temporal patterns of agonistic/defensive behaviors and how we are able to divide the timeline of behaviors into three phases using two different methods to determine the point of social role resolution.

#### Burst detection

**Fig 2** illustrates the occurrence of every aggressive (red) and subordinate (blue) behavior exhibited by each animal from two exemplar dyads across each of the five days of behavioral interaction (see **Supplementary Fig S2** for visualizations across all dyads). It is clear that aggressive and subordinate behaviors are not uniformly distributed as they occur in bursts of activity followed by periods of quiescence. We used Kleinberg’s burst detection algorithm to formally identify time periods containing bursts of aggressive and subordinate behavior (yellow bars in **Fig 2**). The median total number of bursts per dyad over all five days was 20 with an IQR of 11-27 (**Table 2**). The minimum number of bursts for any dyad was 5 and maximum was 79. The frequency of bursts significantly decreased by day (β_days_= −0.31 [−0.38, −0.24]). The duration of individual bursts increased by day (β_days_= 0.08 [0.02, 0.13]). Further, most aggressive and subordinate behaviors occurred inside bursts and the proportion of these behaviors that occurred inside bursts significantly increased over days (β_days_= 0.26 [0.19, 0.32]).

**Fig 2.**
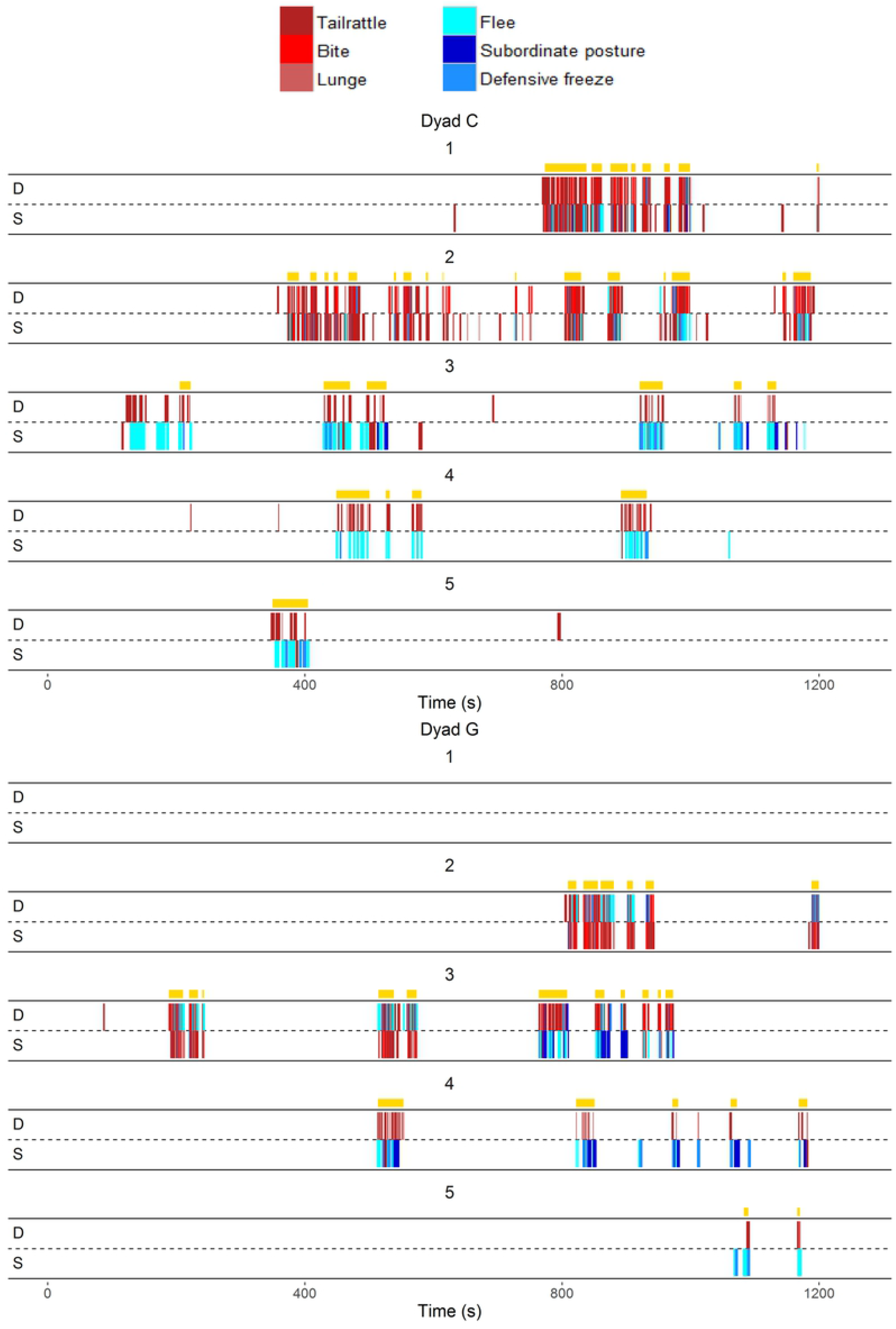
Exemplar temporal pattern of aggressive and subordinate behaviors by the eventual dominant (D) and subordinate (S) males on days 1-5 (Dyad C and Dyad G). Yellow bars represent individual bursts identified by the Kleinberg’s burst detection algorithm (gamma=0.3).

**Table 2.** Descriptive statistics of aggressive and subordinate behavior by day (medians and IQRs)

| Day | Burst frequency | Burst duration (s) | Proportion of aggressive behavior in bursts (%) | Proportion of subordinate behavior in bursts (%) |
| --- | --- | --- | --- | --- |
| 1 | 6 [3, 9] | 14.2 [12.0, 28.9] | 90.8 [83.1, 93.2] | 88.9 [83.4, 95.3] |
| 2 | 3 [2, 10] | 14.4 [11.3, 25.3] | 94.6 [86.3, 97.6] | 94.4 [88.9, 98.9] |
| 3 | 2 [1, 6] | 20.3 [12.2, 39.4] | 91.8 [86.7, 97.0] | 95.3 [91.3, 100.0] |
| 4 | 2 [0, 4] | 19.5 [9.9, 30.0] | 96.5 [88.1, 100.0] | 91.0 [87.3, 98.0] |
| 5 | 1 [1, 4] | 22.1 [12.8, 27.9] | 96.0 [85.7, 99.0] | 98.9 [90.7, 100.0] |
| All | 20 [11, 27] | 15.8 [12.1, 25.3] | 92.0 [88.6, 94.5] | 92.5 [89.2, 95.1] |

#### Identifying resolved dominant-subordinate relationships across bursts

As is evident from **Fig 2** and **Supplementary Fig S2**, both animals in each dyad show a mixture of aggressive and subordinate behaviors during initial interactions. At the end of five days there is one individual that exhibits almost all the aggression without showing subordinate behavior and another individual that shows exclusively subordinate behavior and little aggression. The time point at which this occurs varies by dyad, as does whether this is an abrupt or gradual change. Here, we identified the resolution of each dominant-subordinate relationship (as determined by each animal exhibiting consistent social roles) using two methods, one with very strict criteria (phi-coefficient) and the other with less stringent criteria (using differences in behavior) (see **Methods**).

Phi-coefficients for each burst in each dyad are shown in **Fig 3**. Positive phi-coefficients reflect that the more dominant animal exhibited a higher proportion of aggressive to subordinate behaviors compared to its partner. Relationships were assumed to be resolved at the first burst following which all remaining bursts had ϕ > 0 and p<0.1. Only 8 out of 21 dyads showed a significantly positive phi-coefficient at the first burst and overall phi-coefficients were not significantly different from 0 (Wilcoxon Signed Rank Test, p=0.330) indicating that relationships are typically not resolved during the first aggressive burst. By the final bursts, 19/21 dyads had significantly positive phi-coefficients indicating resolved relationships (binomial test, p<0.001). Of the 2 dyads that did not have significant phi-coefficient at the last burst, one (dyad P) had a phi-coefficient of 1.0 but did not reach significance due to the low frequency of behavior in the burst. The other dyad (dyad R) was the only dyad whose relationship was not resolved by our criteria (see **Methods**) within five days. Using the same criteria, only one relationship (dyad L) was resolved from the very first burst. All 20 resolved dyads reached resolution prior to day five. Four occurred on Day 1, five on each of Days 2 and 3, six on Day 4.

**Fig 3.**
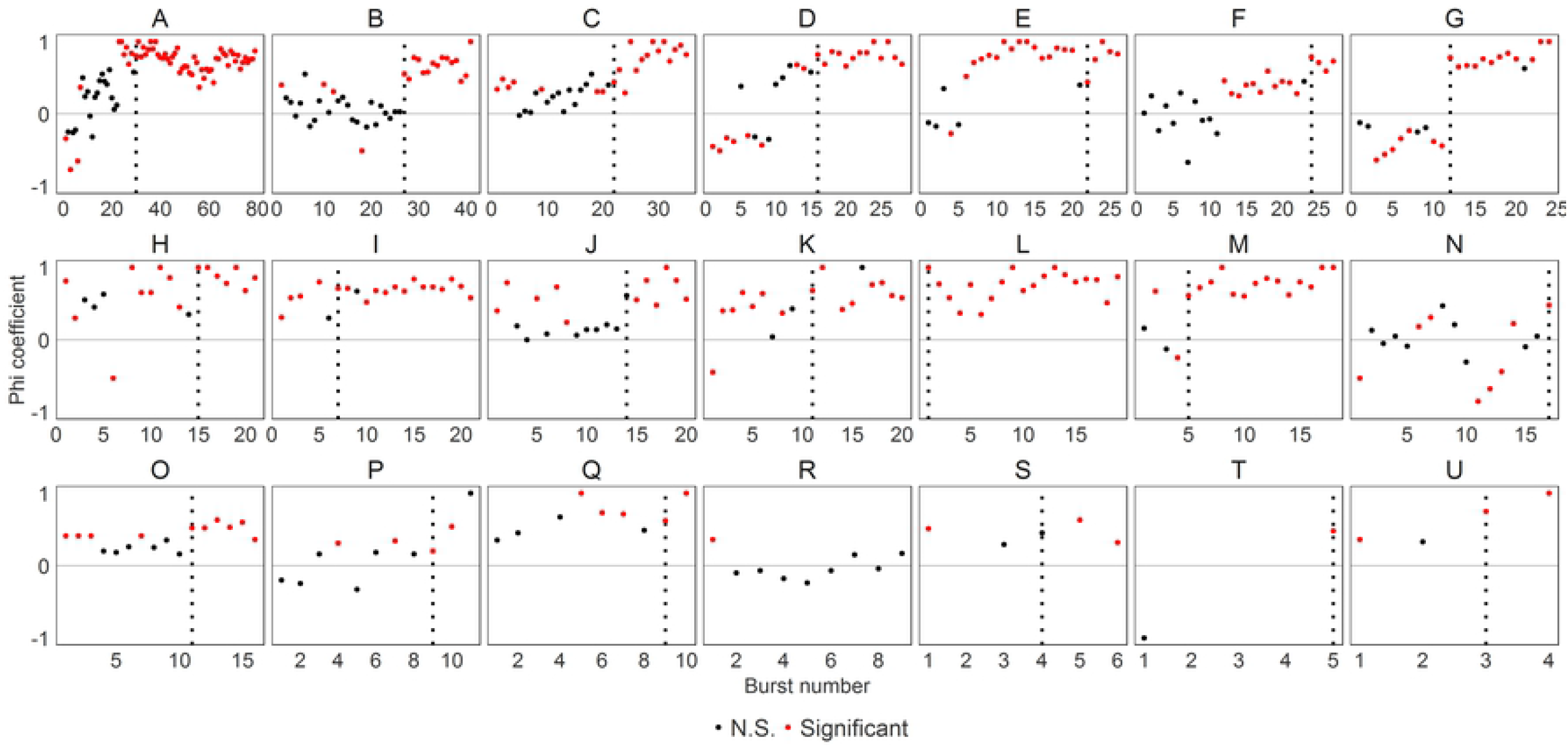
Phi-coefficient of aggressive and subordinate behavior by burst number within each dyad. Black dotted vertical lines represent when relationships are resolved according to phi-coefficient method. Red dots demonstrate significantly distinct levels of aggression versus subordinate behavior by each partner. Note that we excluded any bursts with less than 6 behavior bouts to determine the relationship resolution but those bursts are still shown in the graph.

As shown in **Fig 3**, the phi-coefficient is very stringent, and this may lead to potentially underestimating when relationships become resolved. Therefore we identified the first time point at which dominant-subordinate relationships could potentially be resolved using the difference method. In this method, relationships were considered resolved at the first burst when the difference of aggressive behaviors between the dominant and the subordinate is larger than that of the difference in subordinate behaviors between the subordinate and dominant males for it and all subsequent bursts (see **Methods**). These data are visualized in **Fig 4** – the relationship is resolved at the first burst when the blue line (difference in subordinate behaviors) is below the red line (difference in aggressive behaviors) for all remaining bursts. Using this method, we identified that all relationships were resolved within the observation period. This method identified 13 dyads as having resolved roles on Day 1, three more were resolved on each of Day 2 and 3 and the final two dyads were resolved on Day 4.

**Fig 4.**
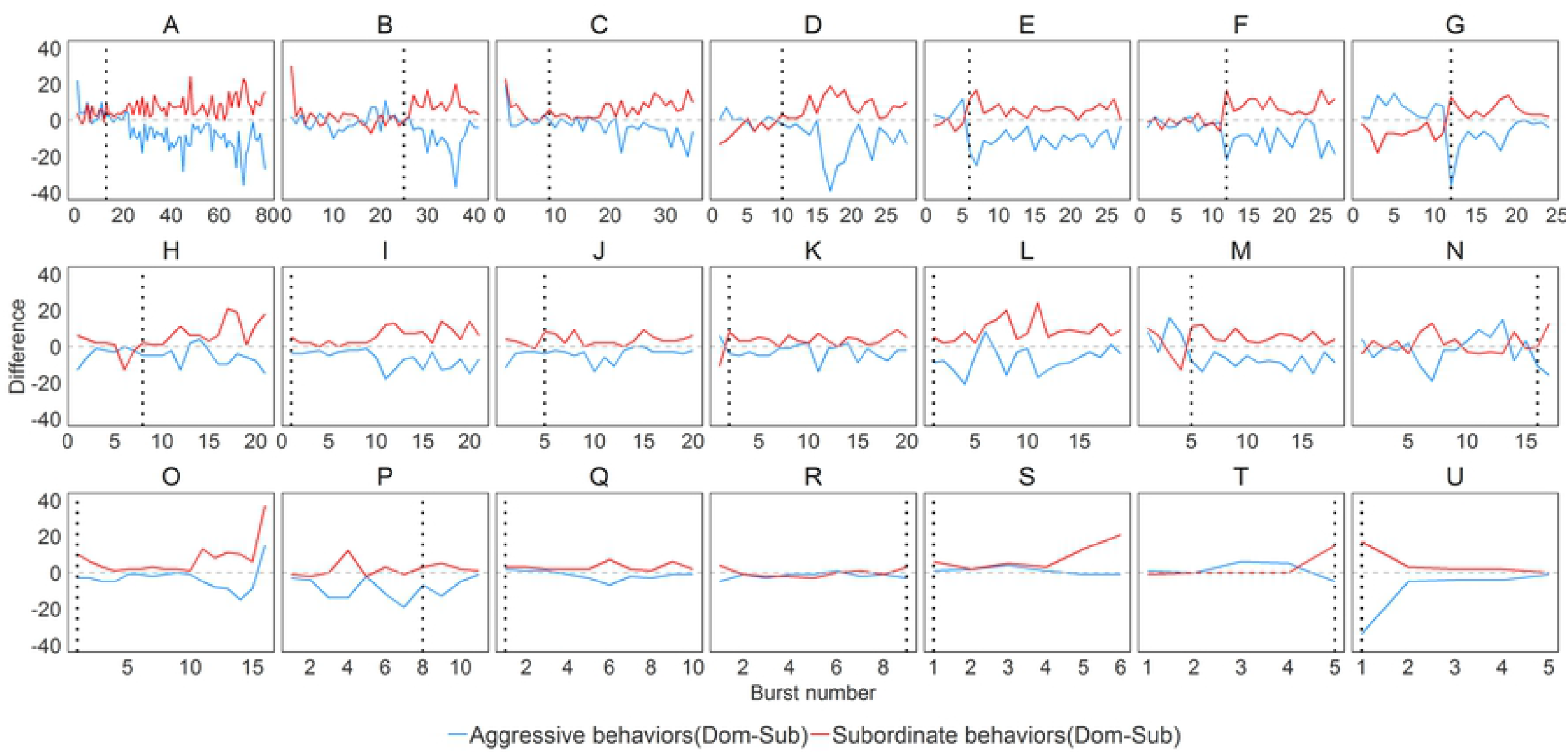
Differences in aggressive and subordinate behaviors between dominant and subordinate males by burst number within each dyad. The red line indicates the difference of aggressive behavior (dominant male – subordinate male) and the blue line indicate difference of subordinate behavior (dominant male – subordinate male). In difference method, the relationship is considered resolved at a point when the blue line is always below red line for all remaining bursts. Black dotted vertical lines represent when relationships are resolved according to the difference method.

The median time to achieve resolution of dominant-subordinate relationship for the phi-coefficient method was after a cumulative 2493 [1502, 3619] seconds of interaction compared to 1054 [629, 2321] seconds for the difference method. By combining the results from these two methods, we were able to identify three separate phases of the relationship development in mice dyads: i) *pre-resolution* – prior to any evidence of social role resolution by either method, ii) *middle* – evidence from the difference method of social role resolution but not fully and consistently resolved according to the phi-coefficient method, iii) *post-resolution (fully resolved)* – significant evidence of each animal exhibiting a consistent social role as confirmed by both methods.

### Behavioral changes across pre-resolution, middle, & post-resolution phases

The changes in the proportions of each behavior exhibited by dominant and subordinate males in each dyad across the three periods are shown in **Fig 5.** Differences in the duration of each behavior between dominant and subordinate individuals at each phase were tested using paired Wilcoxon-Signed Rank Sum Tests. The statistical results are summarized in **Supplementary Table S1.**

**Fig 5.**
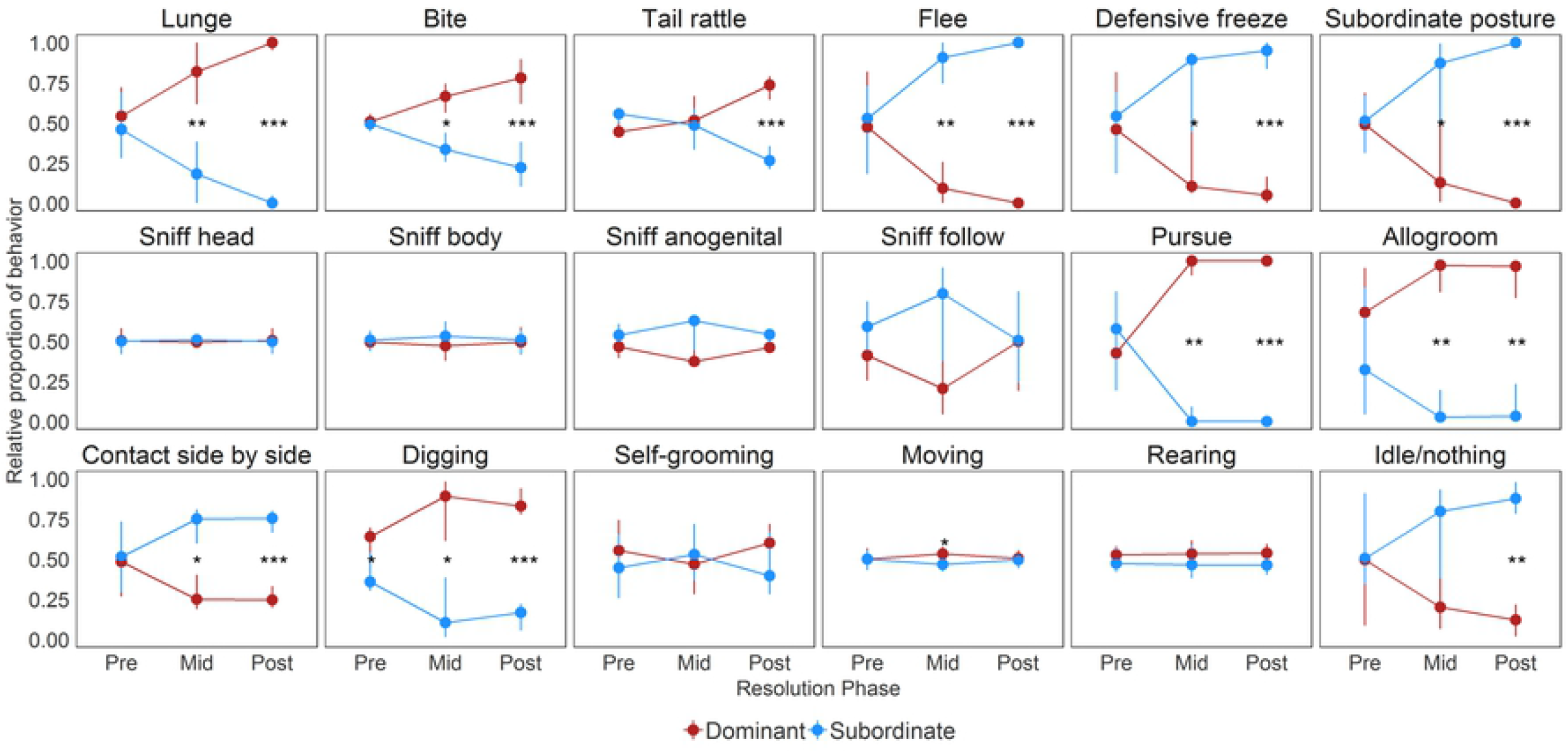
Relative proportion of time each behavior is exhibited the by eventual dominant (red) and subordinate (blue) mice during each phase. Data are medians ± IQRs. The asterisks indicate p-values from paired Wilcoxon Signed Rank Tests; *: p<0.05, **: p<0.01, ***: p<0.001

Unsurprisingly there are clear and consistent changes in aggressive behaviors (lunge, bite, tail rattle) and subordinate behaviors (flee, freeze, subordinate posture) across the three identified phases. Notably, there are no significant differences in aggressive and subordinate behaviors between dominants and subordinates during the pre-resolution phase (all p>0.148). During the middle phase, the differences in these behaviors between dominants and subordinates reached significance (all p<0.041) with the exception of tail rattling (p=0.344). In the post-resolution phase, dominant animals engaged in aggressive behaviors significantly more frequently than subordinates (all p<0.001). Conversely, subordinates spend significantly more time showing subordinate behaviors than dominants (all p<0.001).

Dominants and subordinates differed in the time they engaged in three types of social behaviors (pursue without sniffing, allogrooming, and side-by-side contact) during the middle and post-resolution phases (all p<0.05). In the pre-resolution period there was no significant difference between dominant and subordinate mice in all three of these social behaviors (all p>0.56). During the middle and post-resolution phases, dominants pursued subordinates significantly more (middle: p=0.005, post: p<0.001) and spent significantly longer allogrooming subordinate mice than vice versa (middle: p=0.002, post: p=0.001). Further, subordinates showed significantly more side-by-side contact behavior during the middle (p=0.047) and post-resolution (p<0.001) phases than dominants.

Remarkably, the relative proportion of time spent digging was a strong predictor of dominance. Dominant males exhibited more digging behavior than subordinates in the pre-resolution (p=0.033), the middle (p=0.015) and the post-resolution (p<0.001) phases. Dominants showed more digging than subordinates in 15/21 dyads pre-resolution and 19/21 dyads post-resolution.

Additionally, dominant mice engaged in active moving more frequently than subordinate mice during the middle phase (p=0.047) and subordinates showed more time of inactivity (idle; p=0.001) than dominants during the post-resolution phase. There were no significant differences in the relative proportion of all investigative behaviors (sniff-head, sniff-body, sniff-anogenital, sniff-follow), self-grooming, rearing, and jumping between dominants and subordinates across all three phases (all p>0.07).

### Sequential patterning of behaviors in pre- and post-resolution

We next examined changes in the sequential transition patterns of behavioral states and behaviors from the pre-resolution to post-resolution phases to further examine the microstructure of behavioral sequences.

#### First Order Markov Chains: Event sequences within individuals

Likelihood ratio tests for all individual sequences were significant indicating that behavioral transitions were non-uniformly distributed. We first examined whether dominant and subordinate animals showed difference in transitions from one behavior to another within each individual. In **Fig 6** we highlight all transitions that occurred with a median probability of at least 0.075 (see **Methods**) and color-coded those transitions that occur significantly more frequently than chance in red (also see **Supplementary Table S2**).

**Fig 6.**
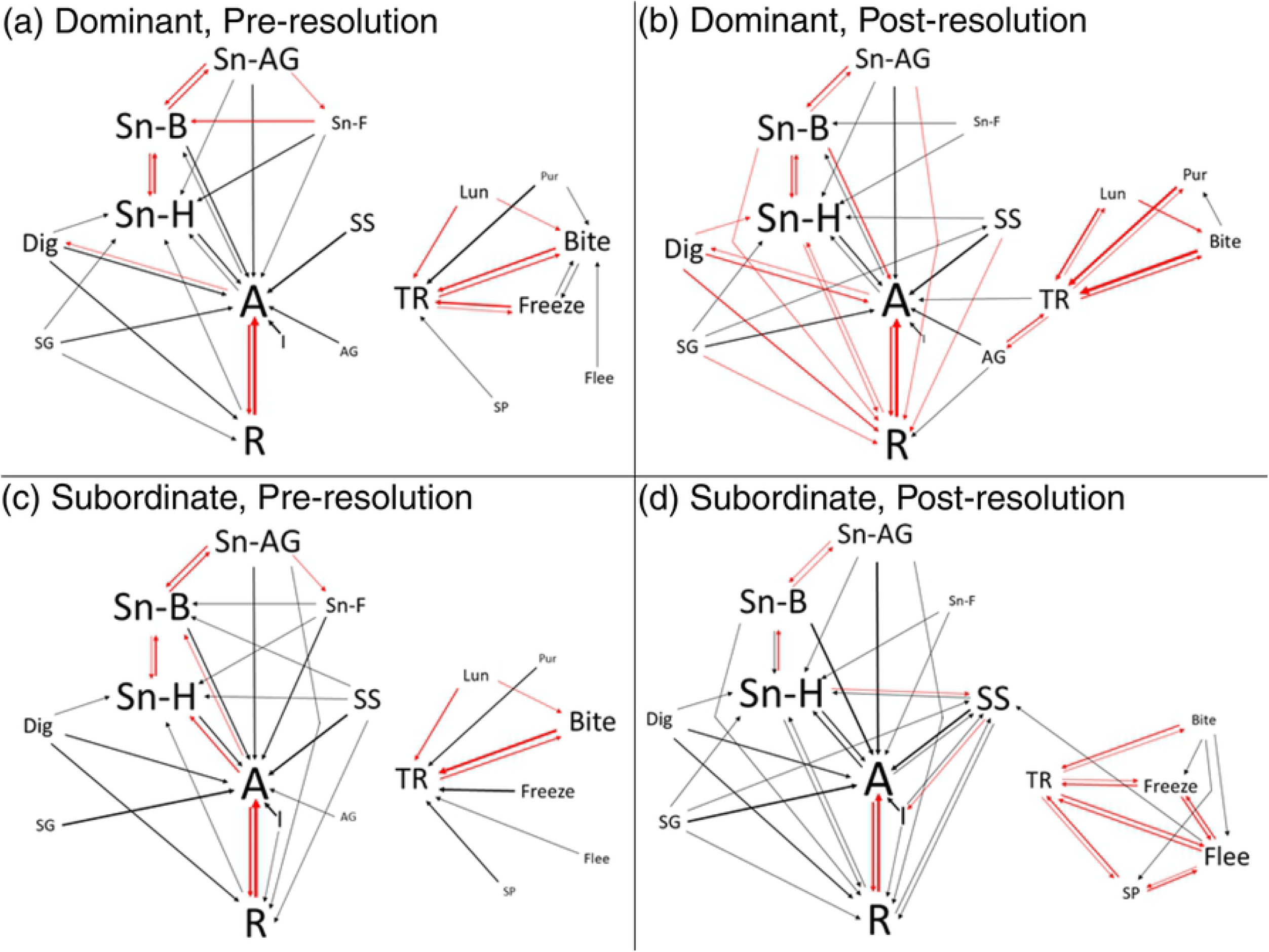
Kinetogram of transitions between behaviors for (A) dominants pre-resolution, (B) dominants post-resolution, (C) subordinates pre-resolution and (D) subordinates post-resolution. All transitions that occur with a median probability of above 0.075 are shown. Label size represents the relative frequency of each behavior. Line weight represents the relative transition probability. Red lines indicate transitions that occur at rates significantly greater than expected by chance. Abbreviations: A – moving, AG – allogrooming, Bite – biting, Dig – digging, Flee – fleeing, Freeze – freezing, I – inactive, Lun – lunge, Pur – pursuing, R – rearing, SG – self-grooming, Sn-AG – anogenital sniffing, Sn-B – body-sniffing, Sn-F – sniff-following, Sn-H – head-sniffing, SP – subordinate posture, SS – side-by-side contact, TR – tail-rattling.

As clearly shown in **Fig 6**, the overall structure of behavior during social interactions is for animals either to be engaged in repeated bouts of general activity and/or sniffing or to be engaged in bouts of aggressive and/or subordinate behaviors. Animals show very infrequent transitions between these two distinct modes of behavior congruent with the above result that aggressive and subordinate behaviors occur in bursts. We found several significant behavioral transitions with a median probability above 0.075 that were present in both dominants and subordinates during both the pre- and post-resolution phases: bidirectional transitions between moving and rearing (all p<0.001), body-sniffing and anogenital-sniffing (all p<0.01), tail rattling and biting (all p<0.001); and unidirectional transitions from head-sniffing to body-sniffing (all p<0.01).

Several transitions were found to only occur significantly differently between post-resolution dominants and subordinates. Post-resolution dominants showed significant bidirectional transitions between tail-rattling and lunging (p<0.001), tail rattling and allogrooming (p<0.001), tail rattling and pursuing (p<0.001) and unidirectional transitions from lunging to biting (p<0.001) that were not observed in subordinates. Interestingly, the significant transition from lunge to bite was also observed in pre-resolution dominants (p=0.002), suggesting this could possibly reflect a difference in fighting strategy that ultimately leads to dominant status. Similarly, pre- and post-resolution dominant mice were the only individuals to show a significant transition from being active to digging (p<0.001). Conversely, post-resolution subordinate mice developed significant bidirectional transitions between tail rattling and all subordinate behaviors (all p<0.01), and between fleeing and freezing (p<0.001) and between fleeing and subordinate posture (p=0.006). These results indicate that established dominant and subordinate individuals develop distinct patterns of behavioral transitions between tail rattling and other aggressive/subordinate behaviors. After relationship resolution, subdominant males still tail rattle but this is less likely to lead to aggressive behaviors and more likely to lead to subordinate behaviors. Further, post-resolution subordinates were also the only group not to show a significant transition from sniffing body to sniffing head, but they did show a significant transition from sniffing head to side-by-side contact (p=0.006).

#### Timed-window cross-correlation between individuals

We used timed-window cross-correlation to determine the likelihood of behavioral states being exhibited by an animal within 2 seconds of when their partner exhibited a specific behavioral state during the pre- and post-resolution phases. In **Fig 7**, we show the FSTTC contingency values between lunging/biting, tail rattling and subordinate behaviors. Based on our finding that tail rattling appeared to act as an indicator of arousal for both dominant and subordinate individuals we chose to examine this behavior separately from the other aggressive behaviors. In **Supplementary Table S3**, we present the FSTTC values of all contingencies and the result of paired Wilcoxon Signed Rank Sum Tests of each contingency between DOM➔SUB and SUB➔DOM directions in both phases.

**Fig 7.**
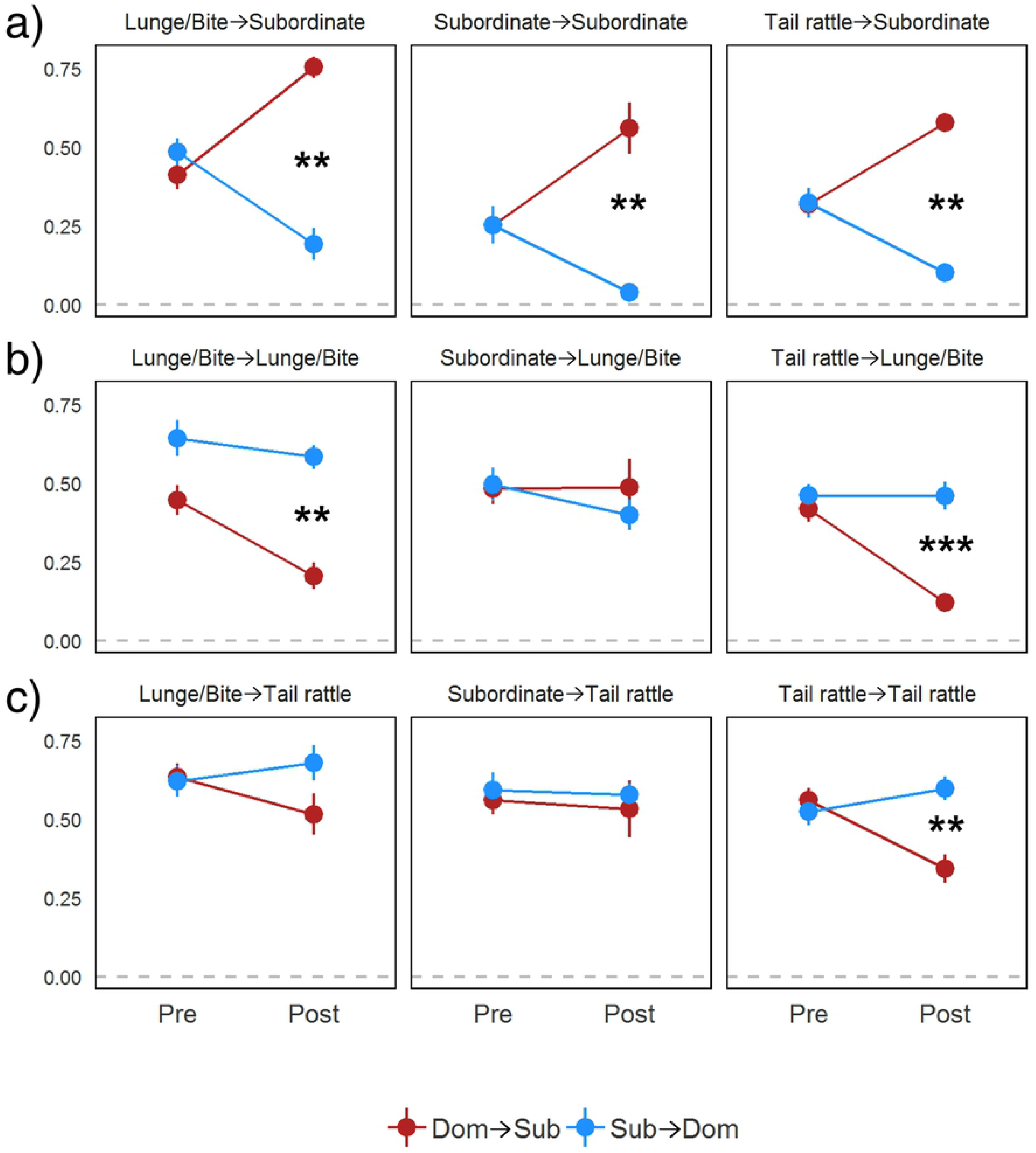
Forward STTC values with standard errors of nine selected contingencies in DOM➔SUB (red) and SUB➔DOM (blue) directions in pre- and post-resolution phases. The asterisks indicate significant differences between DOM➔SUB and SUB➔DOM directions. The differences in FSTTC values between the directions of each dyad were tested using paired Wilcoxon Signed Rank Tests. The asterisks indicate p-values from paired Wilcoxon Signed Rank Tests; *: p<0.05, **: p<0.01, ***: p<0.001

We find strong evidence that subordinate animals learn to consistently yield in response to the behavior of dominant animals but dominant animals become less likely to yield in response to their partner’s behavior as relationships resolve. Pre-resolution, both eventual dominant and subordinate animals show subordinate behavior in response to biting/lunging, tail rattling or subordinate behavior of their partner with moderately high contingency values. After relationship resolution, dominant animals are much less likely to exhibit subordinate behavior in response to biting/lunging, tail rattling or subordinate behavior exhibited by their partner, whereas eventual subordinate animals significantly increase their contingencies showing subordinate behavior almost always in response to each of these behaviors (**Fig 7A**).

We also find evidence that as relationships resolve, dominant animals continue to respond to aggressive behaviors exhibited by their partner with aggression, but subordinate animals decrease this contingency. We found no significant differences in contingency values pre-resolution between dominant and subordinates in biting/lunging or tail rattling leading to biting/lunging. Post-resolution, subordinate animals show significant reductions in these contingencies whereas dominant animals maintain high values (**Fig 7B**). Interestingly, both pre- and post-resolution dominant and subordinate animals were highly likely to respond to subordinate behavior by their partner with biting/lunging or tail rattling, indicating that even subordinate animals may seek to be aggressive if their partner shows any sign of yielding. Further, during the pre- and post-resolution phases both dominant and subordinate animals respond to lunging and biting by their partner with tail rattling suggesting that tail rattling is an indicator of arousal for all individuals. Interestingly, this behavior may actually be used as a signal to deescalate aggressive behavior in pairs. Post-resolution, subordinates significantly decrease their tail rattling in response to the tail rattling by dominant animals whereas dominants continue to show high contingencies of tail rattling in response to their partner’s tail rattling (**Fig 7C**).

## Discussion

In this study, we show that aggressive-subordinate interactions occur in a bursting pattern rather than being evenly distributed across social interactions. We further demonstrate that a series of aggressive-subordinate bursts can be temporally categorized into three phases; pre-relationship resolution, middle, and post-relationship resolution. As animals resolve their dyadic relationship, they significantly change the relative proportion of behaviors they exhibit. Changes are observed not just in aggressive and subordinate behaviors but also in pursuing, allogrooming, and side-by-side contact behaviors. Further, we show that dominant and subordinate mice are very similar in their likelihood to transition from one behavior to the other pre-relationship resolution. However, as they resolve their social relationship, dominant and subordinate animals are significantly different in the frequencies of these transitions. We further demonstrate that animals are highly likely to transition from one behavior to other behaviors within the same behavior states, and that switching from general and social investigation to aggression/subordinate behavior is much less common. We also show that there are temporal changes in how likely dominant and subordinate animals are to rapidly respond to specific behaviors of their partner.

Our analyses indicated that the duration of aggressive/subordinate interactions decreased when animals were exposed to the same individual over five consecutive days. Drews [6] suggests that the hallmark of social dominance is this de-escalation rather than an escalation of aggression once animals establish dyadic dominance relationships or form a social hierarchy in a group. Indeed, many animal species such as lizards [37], fish [38], ants [39], domestic fowls [40] and mice [41] have been shown to decrease the duration or intensity of fights as they establish dominance relationships. Considering each bout of aggressive interaction is energetically costly [42], this decreasing pattern of aggressive interactions suggests that both dominant and subordinate animals benefit from establishing a dominant-subordinate relationship. However, as shown in **Fig 2**, there is variation across dyads in the number of aggressive/subordinate interactions that occur before relationships are resolved. This could be due to variation in the relative differences between partners in competitive ability. Some dyads may be very similarly matched in terms of competitive ability whereas others are very unequally matched. Competitive ability could include many aspects of individual traits such as fitness or energy reserves [43], boldness [44,45], aggression levels [46] or prior experience of winning contests [47]. Alternatively, differences in the time taken to resolve dominance/subordinate status may be reflective of individual differences in social competence. More socially competent individuals may be quicker to establish their social roles and do so with fewer aggressive interactions [3,26].

It is clear that aggressive/subordinate interactions occur in a bursting pattern rather than being distributed evenly during each trial of social exposures. The results of both the First Order Markov Chain and Forward STTC analyses are congruent in demonstrating that transitions within similar behavioral states are most likely and that relatively few behavioral transitions occur across behavioral states. This finding corresponds with previous studies that have shown that groups of mice and chickens show similar bursting patterns of aggressive/subordinate interactions [48,49]. Other groups have looked at mean gap times between bouts of behavior which also show that aggressive social interactions occur in a bursting pattern [50]. Natarajan et al. [16] used First Order Markov Chain analysis to examine strain differences in behavioral transitions during dyadic social interactions and also describe that aggressive behaviors significantly cluster together across strains. At the neurobiological level, this bursting is likely to occur when the neural circuits for aggression become disinhibited and neural circuits for other behaviors (e.g. feeding, repetitive behaviors) are inhibited following exposure to relevant social stimuli [51,52]. For example, Hong et al. [53] showed a glutamatergic neuron subpopulation in the medial amygdala (MeA) promote repetitive self-grooming and concurrently inhibit aggression whereas a GABAergic neuron subpopulation promote aggression while concurrently inhibiting repetitive behaviors. Interestingly, we also observed that as relationships became resolved, a significantly higher proportion of aggressive/subordinate behavior exhibited by both animals occurred within the contexts of bursts indicative of social status stabilizing over time.

During the pre-resolution phase, we found that the eventual dominant and subordinate mice are highly similar in both the frequencies of each behavior and the transition patterns between behaviors. Post-resolution, dominant and subordinate mice diverge significantly in their frequencies and behavioral transition patterns. Unsurprisingly, dominant mice showed higher frequencies of aggressive behavior over all. Lunging was almost exclusively exhibited by dominant animals demonstrating that subordinate mice rarely initiate fights with their more dominant partners. Though subordinates did display biting behavior, about 75% of all bites were exhibited by dominants. Tail rattling was significantly more often observed among dominant compared to subordinate mice, though subordinate mice also showed this behavior post-resolution. Conversely, dominant mice almost never exhibited defensive freezing, subordinate postures or fleeing with all three of these behaviors being highly deterministic of subordination. These findings were confirmed by the FSTTC analysis which found that as relationships resolve, dominant mice continued to show biting and lunging in response to both subordinate and aggressive behavior by subordinate mice, but subordinate mice were much more likely to show subordinate behaviors following biting and lunging by dominant mice. This finding is consistent with the hypothesis that dominant-subordinate relationships are determined when one animal learns to consistently yield to another [6]. We also observed that the behavior ‘pursuing’ was almost exclusive to dominant animals post-resolution. Although this behavior was defined as following without sniffing and is not an overt chase or classically agonistic, it does appear to be diagnostic for dominance as subordinate mice do not ever pursue dominants.

In terms of behavioral transitions within individuals, pre-resolution and post-resolution dominants and subordinates showed large overall similarities in the structural pattern of First Order Markov transitions. This finding suggests that the patterns of behavioral sequences mice express during social interactions are quite stereotyped. The most notable differences observed between groups were for transitions involving tail rattling. We found that post-resolution dominants showed the most frequent transition from tail rattling to lunge or pursuing, whereas post-resolution subordinates showed transitions from tail rattling to all three subordinate behaviors. When comparing behavioral transitions between individuals using the FSTTC analysis we found that both dominants and subordinates are highly likely to show tail rattling in response to their opponents’ aggression in post-resolution. Notably, mice also respond to tail rattling by their partner according to their own social status. Dominants respond to the tail rattling of subordinates with aggressive behaviors while subordinate mice respond to the tail rattling of dominants with subordinate behaviors. Further, subordinates become less likely to respond to the tail rattling of their partners with tail rattling as relationships resolve. This finding suggests that tail rattling may well be an indicator of arousal of both aggressive and defensive neural circuits, with subordinate animals inhibiting the progression of behaviors to aggression following such arousal and instead engaging in subordinate behaviors. Although tail rattling has classically been considered to be an aggressive behavior [54], some studies have also considered tail rattling to be indicative of generalized arousal [54,55] or emotional states [56]. Our findings suggest it is important to consider the social status and context of individuals when interpreting the function of tail rattling.

Post-resolution dominant animals were found to exhibit bidirectional transitions between tail rattling and allogrooming. Further, dominant mice spent significantly longer allogrooming subordinate mice than visa-versa. Several studies have suggested that allogrooming serves an affiliative function promoting positive social relationships [57,58]. Others have argued that allogrooming acts as an antagonistic behavior [57,59,60], which in extreme cases can lead to barbering of subordinates by dominant mice [61–65]. Further, we also found that the frequency of digging behavior was a strong predictor of eventual dominance status. Eventual dominants engaged in this behavior significantly more frequently in both the pre- and post-resolution phases. Indeed, digging was the only behavior pre-resolution that could be used to successfully predict the eventual dominant-subordinate relationship. One of the suggested functions of digging is to enhance scent-marking. Mongolian gerbils show stereotyped digging behaviors while urinating or defecating to preserve the odor of scent-marks for longer [66]. In mice, elevated levels of both digging and grooming behavior have been reported in males who experienced 10 or 20 consecutive days of aggressive victories over subordinate individuals [67,68]. Our results are consistent with these findings and suggest than in pairs with established dominance relationships that both grooming and digging are used by the dominant mouse to reinforce the dominance relationship without requiring them to engage in costly bursts of aggressive behavior. Conversely, we found that post-resolution subordinate mice were more likely to show passive social interaction. Post-resolution dominant mice showed many significant First Order Markov transitions between social behaviors, whereas subordinate mice showed relatively few, consistent with the findings of Koyama *et al*. [69]. Notably, we found that subordinates would not transition from sniffing their partner’s body to sniffing their head suggesting that subordinates are less inclined to actively investigate dominant mice face on. Consistent with this interpretation, we found that post-resolution subordinates showed higher levels of anogenital sniffing which likely occurred when dominants were moving away from them and were unaware of this social contact. We also found that post-resolution subordinates spent longer in side-by-side contact with their partner and were more likely to transition from sniffing their partner’s head to side-by-side contact than dominants. These results appear to be consistent with other data from mice [70] and rats [71] that a loss of social status leads to changes in the pattern of olfactory investigation exhibited by subordinate mice from an active to a passive strategy.

In summary, we show mouse social behaviors can be analyzed in various ways to qualitatively and quantitatively assess the development of a dominant-subordinate relationship in a dyad. We demonstrate that by analyzing the pattern of bursts of aggressive and subordinate behavior it is possible to identify different phases of the social dominance relationships of pairs of mice. These methods may be useful for identifying those animals who are able to form relationships more quickly than others and who may be more socially competent. We also show that examining the First Order Markov transitions between social behaviors within individuals, and the FSTTC contingencies between behaviors of each animal can reveal how patterns of behavior change over time. Developing such methods for analyzing the microstructure of behavior will be helpful for developing our further understanding of the neurobiological mechanisms behind these changes.

## Supporting information

**Supplementary Table S1. The difference in durations of behaviors between dominant and subordinate mice across three phases (paired Wilcoxon Signed Rank Sum Tests).** The value in parenthesis is p-value and asterisks indicate; *: p<0.05, **: p<0.01, ***: p<0.001

**Supplementary Table S2. The median probability of behavioral transition within individuals and its significance tested based on permutation.** Asterisks indicate; **: p<0.01, ***: p<0.001. We conservatively considered the transitions to be significant if p≤0.01 and only significant p-values of transitions are displayed.

**Supplementary Table S3. FSTTC values of each state transitions and the differences in FSTTC values between directions.** The asterisks indicate p-values from paired Wilcoxon Signed Rank Tests; *: p<0.05, **: p<0.01, ***: p<0.001. The values in the parenthesis indicate V value from the tests.

**Supplementary Fig S1. The durations of investigative and social behaviors exhibited by each mouse in each dyad across 5 days of observation.**

**Supplementary Fig S2. Temporal pattern of aggressive (blue) and subordinate (red) behaviors by the eventual dominant (D) and subordinate (S) males on days 1-5 of all 21 dyads.** Yellow bars indicate aggressive/subordinate bursts identified by the Kleinburg Burst Detection algorithm (gamma=0.3).

## Acknowledgements

We thank Dr. Stephen Eglen (University of Cambridge, UK) for his help in writing the permutation algorithm and Dr. Frances Champagne for suggestions in writing the manuscript.

## Author Contributions

Conceived and designed the experiments: JC. Performed the experiments: NB PF JC. Analyzed the data: WL JF JC. Wrote the paper: WL JF JC.

